# Stomatin-like protein 2 senses oxidative stress through the interaction with phosphatidic acid to promote mitochondrial unfolded protein response

**DOI:** 10.1101/2024.04.30.591949

**Authors:** Maria Paulina Castelo Rueda, Irene Pichler, Karolina Musilova, Stanislav Kmoch, Peter P. Pramstaller, Ales Hnizda, Andrew A. Hicks, Roman Vozdek

**Affiliations:** Institute for Biomedicine, Eurac Research, Bolzano, Italy; Research Unit for Rare Diseases, Department of Pediatrics and Inherited Metabolic Disorders, First Faculty of Medicine, Charles University in Prague, Prague, Czech Republic

## Abstract

The mitochondrial unfolded protein response (mtUPR) is an essential mechanism that maintains mitochondrial fitness during stress. Using a genetic screen in *Caenorhabditis elegans* looking for regulators of the mtUPR, we identified *stl-1*, an ortholog of human Stomatin-like protein 2 (SLP-2), as a positive regulator in healthy mitochondria. The loss of STL-1 and SLP-2 results in an impaired mtUPR in *C. elegans* and human cells, respectively. Both *C. elegans* STL-1 and human SLP-2 are proteins located at the inner mitochondrial membrane and exhibit strong lipid binding affinity to phosphatidic acid. Oxidative stress alters the STL-1 localization within the mitochondrial membrane, and triggers the mtUPR dependent on both STL-1/SLP-2 and mitochondrial PA homeostasis. These results reveal an evolutionarily conserved mechanism of mitochondrial protection, in which STL-1/SLP-2 acts as a sensor for changes in mitochondrial membrane lipid composition through physical interaction with PA species, thereby mediating the mtUPR and enhancing stress resistance.

## INTRODUCTION

Mitochondria are vital cellular organelles, in which oxygen is used as the electron acceptor to make energy in the form of ATP through oxidative phosphorylation. Cells are placed in immediate danger when oxidative phosphorylation is blocked, and unconsumed oxygen forms radical agents that cause oxidative stress. Such stressed mitochondrial compartments have intrinsic mechanisms that prevent unbearable damage of the mitochondrial content as well as of the hosting cell and the entire organism, which include mitophagy of the damaged mitochondria, and stimulation of the mitochondrial unfolded protein response (mtUPR). The mtUPR activates expression of nuclear-encoded cytoprotective genes, mostly mitochondrial molecular chaperones, to maintain proteostasis when mitochondrial fitness is impaired and increased mitochondrial proteolysis of misfolded proteins is detected (1,2). How the mtUPR is activated in healthy mitochondria or during stress that is not yet damaging, is currently unknown.

We aimed to identify novel components mediating mitochondrial protection using a transgenic *Caenorhabditis elegans* strain carrying a green fluorescent protein (GFP) reporter of the mtUPR, a pioneering model system that has helped to understand fundamental mechanisms of mtUPR activation triggered by stressed mitochondria (3–9). Our preliminary genetic screen revealed that the *stl-1* gene, an ortholog of Stomatin-like protein 2 (SLP-2), might mediate the mtUPR in non-stressed conditions. SLP-2 is an evolutionary conserved promiscuous scaffolding protein for the stabilization of the phospholipid cardiolipin and several proteins at the inner mitochondrial membrane (10–14) and facilitates stress-induced mitochondrial hyper-fusion in *C. elegans* and mammalian cell models (15,16). However, the mode of action of SLP-2 is poorly understood.

In this paper, we identified *C. elegans* STL-1 and its human ortholog SLP-2 as positive regulators of the mtUPR. We explored their role in the mtUPR under basal and stress conditions, examined phosphatidic acid (PA) binding affinities of purified STL-1 and SLP-2, observed altered expression of STL-1 in mitochondria upon modulated PA/cardiolipin homeostasis, and concluded that STL-1/SLP-2, through direct binding to PA within the mitochondrial membrane, promotes the mtUPR, and thus increases animal survival under oxidative stress.

## RESULTS

### 1. *C. elegans stl-1* promotes mtUPR under basal conditions

To study the mechanisms mediating mitochondrial resilience against oxidative stress, we looked for the genes mediating the mtUPR under basal conditions by using a well-established transgenic *C. elegans* strain carrying the GFP-based reporter of mtUPR, *zcIs9(hsp-60p::GFP)* (17). We performed a preliminary RNAi screen of *C. elegans* genes encoding various mitochondrial proteins and found that RNAi against *stl-1* significantly reduced GFP reporter levels as determined by either GFP fluorescent intensity or the western-blot analysis of animal extracts using anti-GFP antibody (Fig. 1, A to C). These data indicated that the mtUPR pathway is active under non-stress (basal) conditions and suggested that STL-1 could mediate the mtUPR in healthy mitochondria. This observation was surprising because the inactivation of the genes encoding major regulators of mitochondrial dynamics (*phb-2/PHB2, eat-3/OPA1*, *fzo-1/MTF2*) that are known to be stabilised by SLP-2, had the opposite effect, causing strong activation of the mtUPR in *C. elegans* (Fig. 1, A to C) (18). To evaluate a role of STL-1 in the mtUPR pathway, we crossed *zcIs9* animals to those carrying a loss-of-function allele *stl-1(tm1544)*, hereafter referred to as the null allele (-), harbouring a deletion of 555*bp,* and *stl-1* alleles *gk118866* and *gk118865,* that encode STL-1 with the missense mutations R51K and M308I, respectively (Fig. 1D). Both R51 and M308 are highly conserved among SLP-2 orthologs, and their position is presumably located in distinct protein domains based on the predicted 3D structural model generated by AlphaFold (Fig. 1, E to F) (19). We found that all three *stl-1* alleles reduced the levels of GFP protein as well as GFP fluorescence under basal conditions (Fig. 1, A to C, and fig. S1A). Since both missense mutations phenocopied the null allele *tm1544*, we conclude that all three mutated *stl-1* alleles cause loss-of-function of STL-1.

**Fig. 1.**
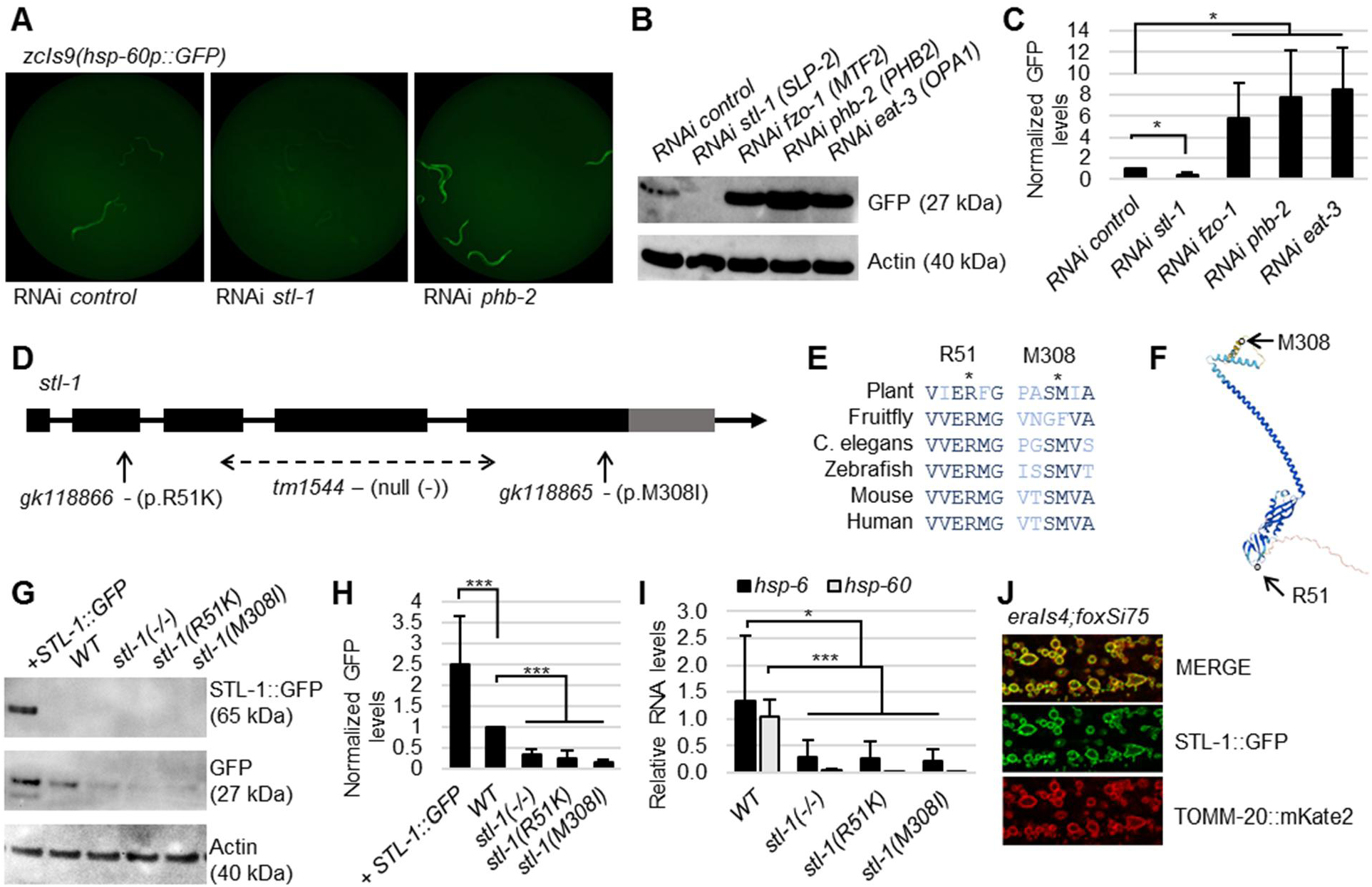
STL-1 promotes the mtUPR under basal conditions. **A.** Widefield fluorescent images of crawling *C. elegans* carrying *zcIs9(hsp-60p::GFP)*, exposed to indicated RNAi condition for three days. **B.** Evaluation of GFP levels by Western blot analysis of ten randomly picked transgenic animals carrying the *zcIs9* transgene and exposed to indicated RNAi. **C.** Semi-quantification of GFP levels from Western blot analysis on the left. n≥ 10 young adult animals for each group with three independent biological replicates, *P< 0.05. **D.** The *stl-1* gene with indicated mutated alleles used in this study. The *tm1544* (null allele (-)) is a 555 *bp* long deletion, *gk118866* is the missense mutation R51K, and *gk118865* is the missense mutation M308I. **E.** Amino acid alignment of SLP-2 orthologs across phyla with indicated amino acids. **F.** Subunit structure of *C. elegans* STL-1 as generated by the AlphaFold and position of amino acid substitutions due to missense mutations. **G.** Evaluation of GFP levels by Western blot analysis of twenty randomly picked animals carrying the *zcIs9* transgene and indicated mutations. **H.** Semi-quantification of GFP levels from Western blot analysis in the indicated strains. n≥ 10 young adult animals for each group with three independent biological replicates, ***P< 0.001. **I.** Relative RNA quantification as determined by RT-PCR of mRNA for *hsp-6* and *hsp-60* extracted from N2 and *stl-1* mutants. Mixed developmental stages for each group with three biological replicates, ***P< 0.001, *P< 0.05. **J.** Fluorescent confocal image of the adult hypoderm. Transgenic animal carries STL-1 reporter *eraIs4(stl-1p::stl-1::GFP::unc-54 3’UTR*) together with the mitochondrial reporter *foxSi75(eft-3p::tomm-20::mKate2::HA::tbb-2 3’UTR)*.

Interestingly, using animals carrying *zcIs13(hsp-6p::GFP)*, another established reporter of mtUPR activation (17), we observed significantly decreased GFP fluorescence only in the *stl-1* mutants with missense mutations R51K and M308I, while the GFP levels in the adult *stl-1* null mutants were not significantly altered compared to wild type (WT) (fig. S1, A to D). These data suggest that the regulation of *hsp-6* expression is distinct from *hsp-60,* and that the *stl-1* null allele could have a detrimental effect on mitochondria, resulting in *stl-1*-independent regulation of selective mtUPR targets. To verify the role of *stl-1* on the endogenous mtUPR target genes, we assessed expression levels of the *hsp-60* and *hsp-6* in *stl-1* mutants using quantitative reverse transcription polymerase chain reaction (qRT-PCR). We found that both *hsp-60* and *hsp-6* mRNA levels were reduced in animals carrying the *stl-1* loss-of-function alleles (Fig. 1I). To further support the role of STL-1 in the mtUPR, we assessed the response in animals with overexpressed STL-1. We generated transgenic animals carrying *eraIs4(stl-1p::stl-1::GFP)*, which harbours several copies of the entire *stl-1* gene together with its 2000 bp upstream sequence, cloned in-frame to the GFP gene. First, we verified that the STL-1::GFP fusion protein is expressed and correctly localised to the mitochondrial membrane, marked by the TOMM-20::mKate2 reporter (Fig. 1J). We subsequently crossed *zcIs9 with eraIs4* and found that expressed STL-1::GFP significantly activated the *zcIs9(hsp-60p::GFP)* transgene (Fig. 1, A to C). These data together indicate that mitochondrial STL-1 promotes expression of the mtUPR target genes under basal conditions.

### 2. The *stl-1* gene mediates mtUPR upon oxidative stress

We hypothesized that STL-1 could protect mitochondria from oxidative stress by stimulating mitochondrial protective mechanisms through activation of the mtUPR. We exposed WT and *stl-1* mutants carrying *zcIs9* to 100µM paraquat, an exogenous toxin, which upregulates the mtUPR, disrupts mitochondrial membrane permeability, and reduces activity of mitochondrial complexes I, III, and IV (20). We found that the increased levels of GFP reporter upon paraquat exposure were significantly suppressed in all three *stl-1* mutants carrying *zcIs9(hsp-60p::GFP)* compared to the parental strain (Fig. 2, A to C). Then, we evaluated expression of endogenous *hsp-60* and *hsp-6* mRNA levels after exposure to paraquat and found a different effect between GFP reporter levels and mRNA levels of the endogenous mtUPR target genes among tested *stl-1* alleles. Specifically, we determined that *hsp-60* mRNA levels were suppressed in *stl-1(-/-)* and *stl-1(M308I)* mutants but not in *stl-1(R51K)* mutants, while the levels of *hsp-6* mRNA were significantly altered in *stl-1(R51K)* mutants, but not in *stl-1(-/-)* and *stl-1(M308I)* mutants (fig. S2A). These data further support the fact that different mtUPR target genes may have distinct activation pathways, and that various *stl-1* mutants exhibit different dynamics of mtUPR regulation.

**Fig. 2.**
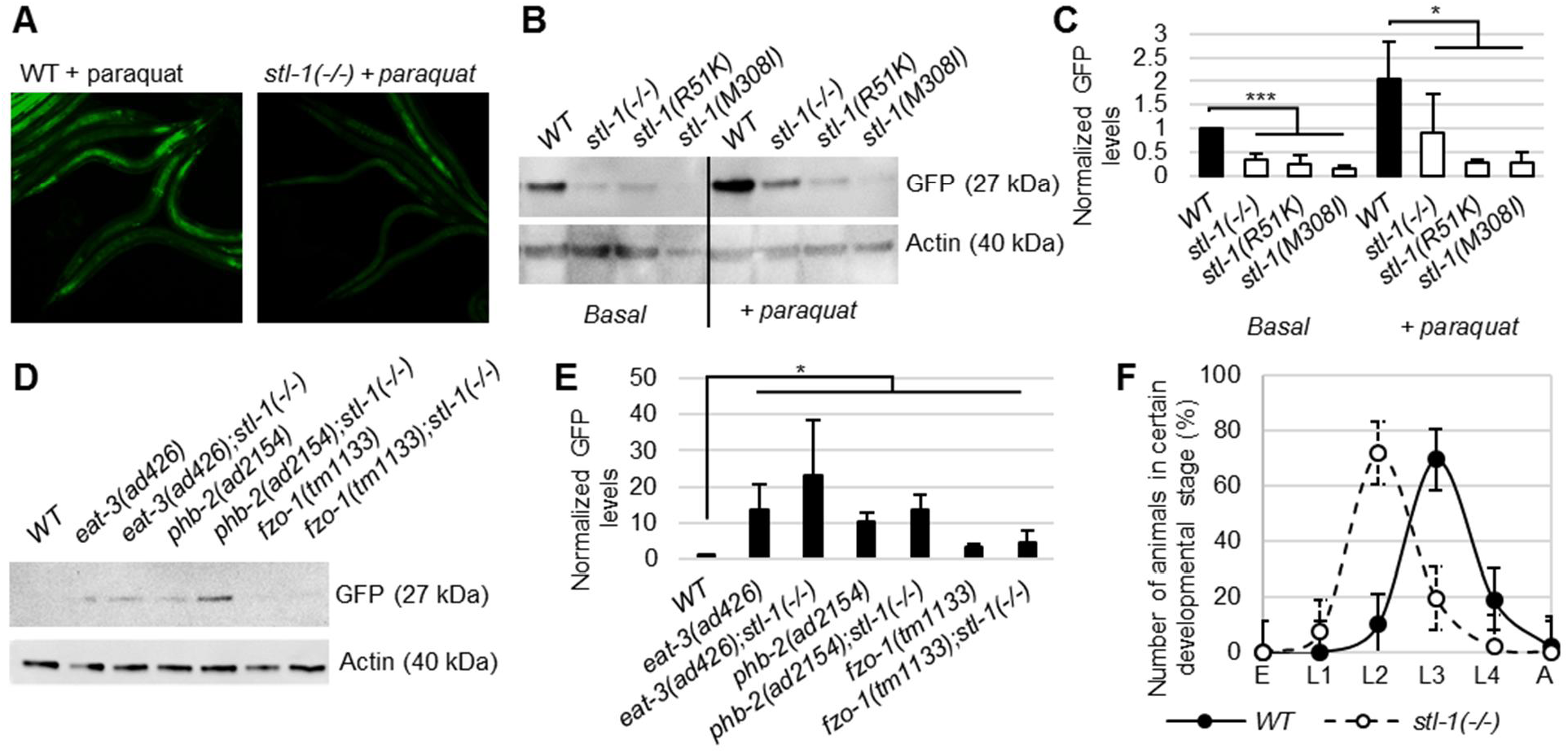
The role of STL-1 in the mtUPR upon different stress conditions. **A.** Widefield fluorescent images of *C. elegans* carrying *zcIs9* after exposure to paraquat for three days. **B.** Evaluation of GFP levels by Western blot analysis of ten randomly picked adult animals carrying the *zcIs9* transgene, indicated allele and grown in specified condition. **C.** Semi-quantification of GFP levels from Western blot analysis on the left. n ≥ 10 young adult animals for each group with three independent biological replicates, ***P< 0.001, *P < 0.05. **D.** Evaluation of GFP levels by Western blot analysis of ten randomly picked transgenic animals carrying the *zcIs9* transgene and indicated mutated alleles. **E.** Semi-quantification of GFP levels from Western blot analysis on the left. n≥ 10 young adult animals for each group with three independent biological replicates, *P < 0.05. **F.** Distribution of developmental stages of the *C. elegans* WT and *stl-1(-/-)* mutants after 3 days of development in the presence of paraquat. n ≥ 20 animals for each group with three independent biological replicates, ***P< 0.001.

The fact that the GFP reporter levels were substantially increased in the *stl-1* mutants after exposure to paraquat as compared to basal conditions indicates that the mtUPR is activated in both an *stl-1*-dependent and *stl-1*-independent manner. Therefore, we tested whether *stl-1* is required for mtUPR activation when mitochondrial dynamics are impaired. We used loss of function alleles *phb-2(ad2154)* (21), *eat-3(ad426)* (22) and *fzo-1(tm1133)* (23) causing mitochondrial fragmentation, and verified that mutated *phb-2, eat-3* as well as *fzo-1* robustly increased GFP levels in *zcIs9* animals, thus resembling the activated mtUPR phenotype, upon their inactivation by RNAi (Fig. 1, B and C, and Fig. 2, D and E). We subsequently crossed *phb-2, eat-3* and *fzo-1* mutants with *stl-1* mutants and observed that the GFP levels were further amplified in double knockouts (Fig. 2, D and E, and fig. S2, B and C). These data indicate that genetically compromised mitochondrial fusion leads to mtUPR activation independently of *stl-1* and suggests that loss of *stl-1* in *phb-2*, *eat-3* and *fzo-1* mutants may further compromise mitochondrial fitness that triggers robust mtUPR activation.

Since the *stl-1* mutants are superficially wildtype under standard conditions and develop within three days to adulthood, we determined whether the suppressed mtUPR in *stl-1* mutants might have a beneficial or a detrimental effect on animal development upon oxidative stress. We evaluated animal development during exposure to 100µM paraquat, the concentration that activates the mtUPR in WT but not in *stl-1* mutants, and found that *stl-1* knockout animals showed delayed development. Specifically, within five days, most *stl-1* mutants developed into the L2 larval stage compared to WT animals, which developed into the L3-L4 larval stages (Fig. 2F). These data demonstrate a critical role of *stl-1* under exogenous oxidative stress. Based on the data collected, we conclude that STL-1 stimulates mitochondrial protective mechanisms in non-stressed animals causing a mitohormesis-like state, ensuring sufficient levels of mtUPR targets in the cells to increase the threshold to toxic stimuli, while in compromised mitochondria, the role of STL-1 in the mtUPR is diminished and the mtUPR is activated more through *stl-1*-independent pathways.

### 3. STL-1 exhibits pH-dependent binding to phosphatidic acid

To understand the underlying mechanisms of the *stl-1*-mediated mtUPR in healthy mitochondria, we evaluated the biochemical properties of the STL-1 protein. STL-1/SLP-2 belongs to the SPFH protein family characterized by membrane-associated high-order oligomeric assemblies (24). Previous studies have shown that human SLP-2 is a membrane-associated protein localized in the inner mitochondrial membrane, where it can bind to cardiolipin (11). To explore the lipid binding activity of STL-1 protein, we captured STL-1::GFP from *eraIs4* animals using GFP affinity chromatography (fig. S3A) and subsequently used purified STL-1::GFP in a lipid overlay assay. We tested lipid binding activity using a lipid membrane strip (Fig. 3A) and found that purified STL-1::GFP predominantly binds to phosphatidic acid (PA) and, to a lesser extent, to phosphatidylinositol (3,4,5)-trisphosphate (PIP3), phosphatidylinositol (4,5)-bisphosphate (PIP2), and phosphatidylinositol (4)-phosphate (PIP) (Fig. 3, B and C).

**Fig. 3.**
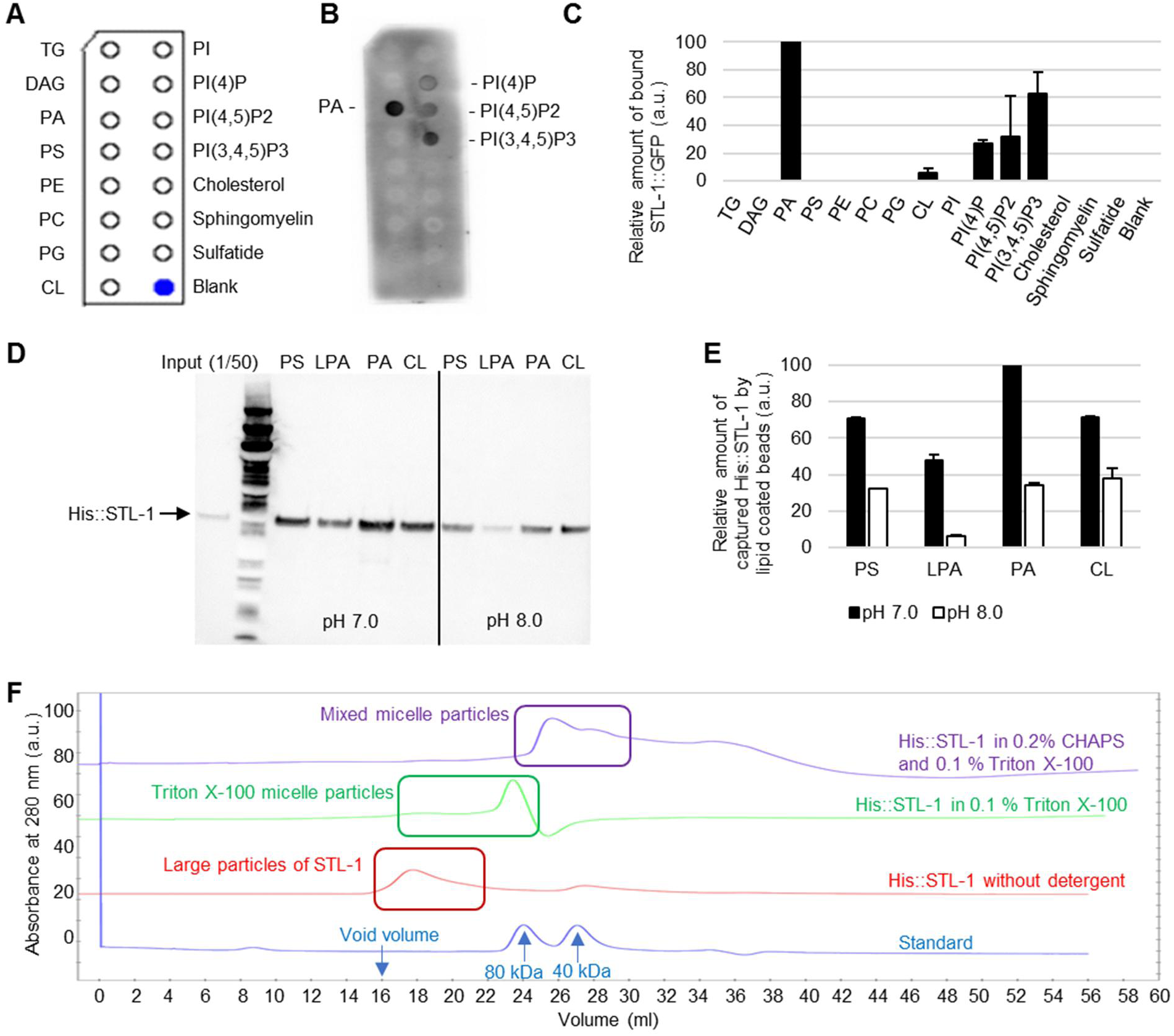
Purified STL-1 exhibits highest binding affinity to phosphatidic acid. **A.** Design of the Echelon membrane lipid strip. TG; Triglyceride, DAG; Diacylglycerol, PA; Phosphatidic acid, PS; Phosphatidylserine, PE; Phosphatidylethanolamine, PC; Phosphatidylcholine, PG; Phosphatidylglycerol, CL; Cardiolipin, PI; Phosphatidylinositol. **B.** Lipid overlay assay using purified STL-1::GFP and visualized by anti-GFP antibody. **C.** Semi-quantification of bound STL-1::GFP on indicated lipid spots with three technical replicates. **D.** Pull-downs using lipid-coated beads and purified His::STL-1 analysed by SDS-PAGE followed by western blot analysis using anti-His-tag. Specific lipids coated on the beads are as follows: PS; Phosphatidylserine, LPA; Lysophosphatidic acid, PA; Phosphatidic acid or CL; Cardiolipin. **E.** Semi-quantification of pulled down His::STL-1 by specified lipid beads with three technical replicates. **F.** Size exclusion chromatography elution profiles of His::STL-1 proteins in buffers with varying composition of detergents. Highlighted areas indicate fractions containing His::STL-1 as determined by SDS-PAGE. a.u.; arbitrary units.

Given that the STL-1::GFP purified directly from *C. elegans* tissues may be in a complex with its intrinsic interacting partners, or that the GFP tag could interfere with the lipid binding, we further evaluated the lipid binding specificity of STL-1 using bacterially expressed recombinant STL-1. The recombinant STL-1 containing an N-terminal 6xHis-tag replacing its mitochondrial signal sequence, was purified using Nickel resin (fig. 3, B and C). Purified His::STL-1 was used in the same lipid overlay assay and was also found to bind to PA and sulfatide (fig. S3D). Next, we explored whether STL-1 can specifically bind PA in solution, using protein pull-down by lipid-coated beads. We assessed binding to beads carrying various membrane lipids, including PA, cardiolipin (CL), Phosphatidylserine (PS), and Lysophosphatidic acid (LPA) at pH 7.0 and pH 8.0. First, we determined that recombinant His::STL-1 binds to all tested lipid beads. The beads carrying PA, however, captured the highest amount of the His::STL-1, further supporting the high affinity STL-1 binding to PA (Fig. 3D). Second, we found that lipid binding of His::STL-1 decreased at pH 8.0 using all tested lipid beads, indicating that pH could be a regulatory factor for STL-1 interaction with phospholipids (Fig. 3, D and E).

Intriguingly, we observed that concentration and physico-chemical properties of detergents used during the purification procedure significantly altered the size of STL-1 particles obtained (Fig. 3E). Size exclusion chromatography (SEC) in the presence of Triton X-100 revealed an STL-1 elution volume corresponding to about 90 kDa, a size close to that of Triton X-100 micelles. On the other hand, SEC in the presence of CHAPS detergent, which forms considerably smaller micelles of 6 kDa, revealed that His::STL-1was eluted in the void volume which indicates the formation of large self-associated particles, and thus STL-1 failed to be incorporated into CHAPS micelles (fig. S3, D and E). Similar data were obtained when SEC was performed without detergent. Lastly, we observed that addition of Triton X-100 to the buffer without detergent, or replacing CHAPS by Triton X-100, can fully transform the STL-1 aggregates into species whose size corresponds to that of the detergent micelles present in the solution. This demonstrates that aggregation of STL-1 can be reverted by protein incorporation into lipid/detergent particles with proper physical characteristics, especially the size of the micelle (Fig. 3E). These data indicate that STL-1 can bind several lipids, and its binding is dependent on spatial configuration of formed lipid particles. Therefore, we conclude that the pH-dependent interaction between STL-1 and PA is a regulatory factor of STL-1 activity rather than an essential element for STL-1 association with the membrane.

### 4. Mitochondrial phosphatidic acid-cardiolipin homeostasis regulates mtUPR through *stl-1*

To explore the biological implication of STL-1 binding to PA, we studied the role of its metabolism in the mtUPR. Mitochondrial PA serves as a precursor lipid for the CL biosynthesis pathway, which when disrupted leads to compromised mitochondria (25). We used an RNAi screen of genes responsible for the CL biosynthesis machinery to better understand the connection of PA and CL in mtUPR (Fig. 4A). First, we determined that RNAi against the genes of the CL biosynthetic pathway, including Phosphatidylglycerophosphate Synthase *pgs-1*, Cardiolipin Synthase *crls-1,* and Tafazzin *acl-3,* all activated the mtUPR, as observed by the increased GFP fluorescence in knockdowns carrying the *zcIs9(hsp-60p::GFP)* reporter (Fig. 4, B and C, and fig. S4, A and B). Next, we tested the genes responsible for the intermembrane transport of phospholipids, such as PRELIDs (Protein of Relevant Evolutionary and Lymphoid Interest Domain containing proteins) and their interaction partner TRIAP1 (TP53 Regulated Inhibitor Of Apoptosis 1) (26,27). There are two conserved genes in the *C. elegans* genome that are orthologous to human PRELID proteins. The *prel-1* gene encodes the ortholog of PRELID1 responsible for intermembrane transport of PA, while *prel-3* encodes the ortholog of PRELID3A and PRELID3B, responsible for intermembrane transport of PS (fig. S4C). We observed that RNAi against *prel-3*/PRELID3 as well as *mdmh-35*/TRIAP1 strongly induced the GFP fluorescence in *zcIs9* animals, suggesting that disruption of PS transport induces the mtUPR. On the other hand, RNAi against *prel-1*/PRELID1 reduced the GFP reporter levels suggesting that disrupted mitochondrial intermembrane transport of PA suppresses the mtUPR (fig. S4, A and B). To evaluate the CL homeostasis in tested knockdowns, we exposed the animals expressing the mitochondrial reporter TOMM-20::mKate2 to 10-N-nonyl acridine orange (NAO), which exhibits strong colocalization with the mitochondrial membrane due to its high specificity to bind CL (28). Using high-resolution confocal microscopy, we found strong colocalization of NAO with mitochondrially localized TOMM-20::mKate2 in *prel-1*, *pgs-1* and *acl-3* knockdowns, while the mitochondrial colocalization of NAO was disrupted in the *crls-1* knockdowns (Fig. 4, D and E, and fig. S4C). In addition, we observed substantially reduced mitochondrial colocalization of NAO in *prel-3 and mdmh-35* knockdowns, which suggests that *prel-3* together with *mdmh-35* may play a role in CL homeostasis via regulated intermembrane transport of PS. In fact, the mitochondrial membrane in *pgs-1* knockdowns was strongly stained by NAO, indicating that either RNAi against *pgs-1* is not sufficient to completely knock-down *pgs-1* activity to reduce CL levels or that there are alternative pathways to preserve/biosynthesize CL omitting *pgs-1* activity, such as the CL biosynthetic pathway from PS, phosphatidylethanolamine and phosphatidylglycerol in bacteria (29).

**Fig. 4.**
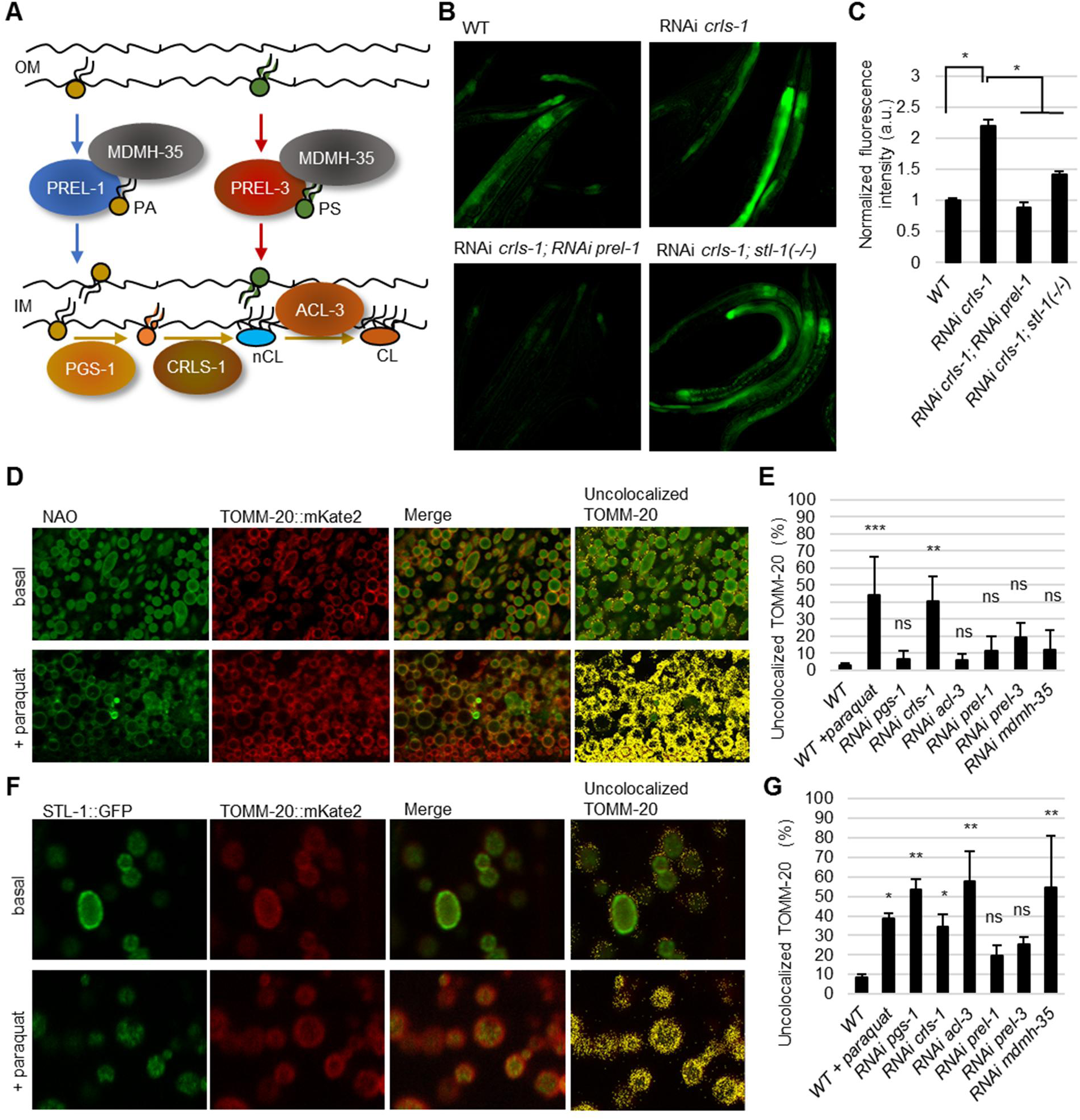
Oxidative stress alters the STL-1::GFP pattern and mitochondrial membrane lipid composition. **A**. Illustration of the roles of specific *C. elegans* orthologs in the mitochondrial phosphatidic acid (PA) and cardiolipin (CL) metabolism. OM; outer membrane, IM; inner membrane, PS; Phosphatidylserine. **B.** Widefield fluorescent images of *C. elegans* carrying *zcIs9* after three days of exposure to specified RNAi condition. **C.** Semi-quantification of fluorescence intensity in *zcIs9* animals and indicated RNAi condition. n ≥ 5 young adult animals per condition, *P < 0.01. ns; not significant. a.u.; arbitrary units. **D.** Confocal fluorescent images of hypodermal tissue in animals carrying *foxSi75* (TOMM-20::mKate2) and grown under basal condition or in the presence of paraquat. Animals were exposed to 10-N-nonyl acridine orange (NAO) for 1 hour before being imaged. **E.** Quantification of TOMM-20::mKate2 signal that does not colocalize with NAO. N ≥ 3 young adult animals per condition, ***P < 0.001, **P < 0.01. ns; not significant. **F.** Confocal fluorescent images of hypodermal tissue in animals *carrying *eraIs4* (STL-1::GFP); *foxSi75* (TOMM-20::mKate2) and grown under basal conditions or in the presence of paraquat. **G.** Quantification of TOMM-20::mKate2 signal that does not colocalize with STL-1::GFP. n ≥ 3 young adult animals per condition, **P < 0.01, *P < 0.05. ns; not significant.

Based on the obtained data we hypothesized that PREL-1 promotes the mtUPR by transporting PA into the inner mitochondrial membrane, and that reduced CL biosynthesis alters the lipid composition of the inner membrane in a way that activates the mtUPR. To define the genetic relationship of *prel-1* to *crls-1*, we performed epistasis analysis by producing double knockdowns elicited by RNAi against *prel-1* and *crls-1*. We found that *prel-1* is epistatic to *crls-1* since inactivated *prel-1* suppressed the mtUPR induced upon *crls-1* inactivation (Fig. 4, A to C). These data suggest a pathway, in which intermembrane transport of PA mediated by PREL-1 promotes mtUPR when CL levels are reduced.

We then examined whether mitochondrial PA homeostasis at the inner mitochondrial membrane influences STL-1 to mediate the mtUPR. First, we evaluated the role of STL-1 in the mtUPR triggered by disrupted CL biosynthesis. We found that the *stl-1* mutants carrying *zcIs9(hsp-60p::GFP)* exhibited a suppressed mtUPR activation upon RNAi against all tested genes, including *pgs-1*, *crls-1*, *acl-3, mdmh-35,* and *prel-3* (Fig. 4, A to C, and fig. S4, A and B). Second, we explored the expression pattern of STL-1::GFP upon modulated mitochondrial membrane lipid composition. We subjected animals carrying *eraIs4(stl-1p::stl-1::GFP)* to RNAi against genes involved in PA/CL metabolism and found that disrupted PA/CL homeostasis resulted in increased formation of clustered STL-1::GFP foci at the inner mitochondrial membrane, characterized by increased levels of TOMM-20::mKate2, which did not colocalize with STL-1::GFP (Fig. 4, F and G, and fig. S4C). Because the expression pattern of STL-1::GFP was significantly altered in *pgs-1*, *crls-1* and *acl-3* knockdowns, we conclude that reduced CL levels at the inner mitochondrial membrane alter the spatial organization of STL-1::GFP. Third, we evaluated the effect of oxidative stress induced by a non-lethal concentration of paraquat on mitochondrial membrane CL levels by NAO staining and the STL-1::GFP expression pattern. We observed that such paraquat treatment resulted in reduced NAO colocalization with the mitochondria, and increased the formation of STL-1::GFP clustered foci that did not colocalize with TOMM-20::mKate2 (Fig. 4, A to C), thus recapitulating phenotypes observed upon inactivated CL biosynthesis. In summary, these data indicate that an altered mitochondrial PA/CL ratio, induced by either genetic intervention or by oxidative stress, alters the expression pattern of STL-1::GFP at the mitochondrial membrane and triggers the mtUPR in an *stl-1*- and *prel-1*-dependent manner.

### 5. The *ymel-1* and *haf-1* mutants resemble mtUPR regulation in *stl-1* mutants

To investigate the possible downstream mechanism of mtUPR activation by STL-1, we explored the role of *C. elegans ymel-1* in the mtUPR. This gene is an ortholog to the human *YME1L1,* which plays a role in protein processing at the inner mitochondrial membrane and can be regulated by SLP-2 (13). We crossed *zcIs9(hsp-60p::GFP)* animals to a loss of function allele of *ymel-1(tm1920)*, which defines a deletion of 759 *bp* over two exons, and found a suppressed mtUPR that phenocopies the *stl-1* null allele (Fig. 5, A to C). Notably, we found a similar reduction of the mtUPR reporter in animals carrying the *haf-1* deletion allele, which was previously identified to suppress the mtUPR (30). The *haf-1* gene encodes the ortholog of the mitochondrial membrane peptide transporter ABCB10 (ATP-binding Cassette, Subfamily B, Member 10), which promotes the mtUPR through the efflux of the protein peptides (30). To investigate whether HAF-1 and/or YMEL-1 could cooperate with STL-1 in mtUPR activation, we evaluated mtUPR reporter levels in *stl-1(-/-); ymel-1(-/-),* and *stl-1(-/-); haf-1(-/-)* double mutants, and found that the GFP levels were not further suppressed as compared with the single mutants (Fig. 5, A to C). These data suggest that *stl-1* might promote the mtUPR through the same signalling pathway as *ymel-1* and/or *haf-1* (Fig. 5, A to C). Next, we compared the mtUPR activation in *ymel-1* and *haf-1* mutants upon paraquat exposure, as well as disrupted CL homeostasis in *crls-1* knockdowns. Using *zcIs9* animals, we found a similar suppression of the GFP reporter levels in *crls-1* knockdowns carrying either *stl-1*(**-/-**), *haf-1(-/-)* or *ymel-1(-/-)* null alleles compared to WT (Fig. 5, E and F), and paraquat-induced mtUPR was similarly suppressed in *stl-1(-/-)* and *haf-1(-/-)* mutants (Fig. 5, A and B). On the other hand, we observed that *ymel-1(-/-)* mutants did not exhibit a suppressed mtUPR upon paraquat exposure. In addition, we found that the animals carrying the *zcIs13(hsp-6p::GFP)* reporter did not show suppressed levels of GFP in any of the mutants carrying a null allele, yet the *ymel-1* mutants exhibited increased GFP fluorescence (fig. S5, A and B). These results further highlight the distinct regulation among different mtUPR targets, and suggest that the *zcIs13(hsp-6p::GFP)* reporter is rather activated in an *stl-1* independent manner. These data together support a model in which STL-1/SLP-2 cooperates with YMEL-1/YME1L and/or HAF-1/ABCB10 at the inner mitochondrial membrane to promote mtUPR in healthy mitochondria.

**Fig. 5.**
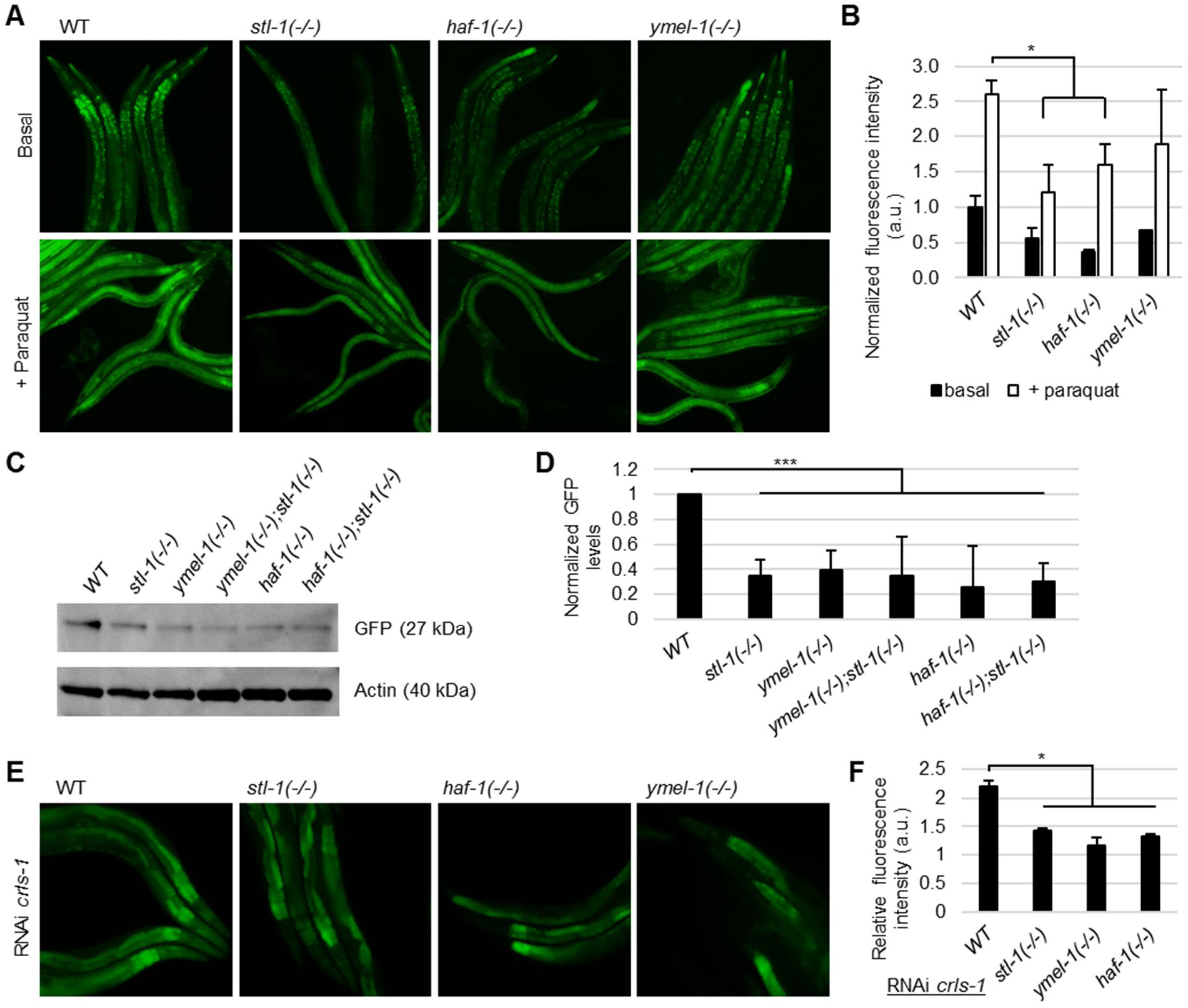
Role of YMEL-1 and HAF-1 in STL-1-mediated mtUPR. **A.** Widefield fluorescent images of indicated *C. elegans* mutants carrying *zcIs9* under basal condition and after three days of exposure to paraquat. **B.** Semi-quantification of fluorescence intensity. n ≥ 5 young adult animals per condition, *P < 0.05. ns; not significant. a.u.; arbitrary units. **C.** Evaluation of GFP levels by Western blot analysis of 10 randomly picked transgenic animals carrying the *zcIs9* transgene and indicated mutated alleles. **D.** Semi-quantification of GFP levels from Western blot analysis. n ≥ 10 young adult animals for each group with three independent biological replicates, ***P < 0.001. **E.** Widefield fluorescent images of indicated *C. elegans* mutants carrying *zcIs9* under RNAi against *crls-1*. **F.** Semi-quantification of fluorescence intensity. n ≥ 5 young adult animals per condition, *P < 0.05. ns; not significant. a.u.; arbitrary units.

### 6. Human SLP-2 binds phosphatidic acid and mediates mtUPR in human cell models

To evaluate the evolutionary conservation of STL-1 regulating the mtUPR, we used SH-SY5Y neuroblastoma cells as a human cellular model system, in which we examined the mtUPR upon basal and stress conditions when SLP-2 was downregulated by expressing a lentivirally transduced shRNA antisense construct for SLP-2. Induction of the mtUPR was determined by the levels of the heat shock proteins GRP75 and HSP-60 (31). Upon 24 hours of exposure to paraquat, we observed increased levels of HSP60, GRP75 as well as SLP-2. However, the cells with a stable SLP-2 knockdown, resulting in a downregulation of about 50% compared to wild type cells, showed significantly decreased levels of both HSP60 and GRP75 under basal conditions as well as following paraquat treatment (Fig. 6, A and B). These data indicate that SLP-2 increases the levels of HSP-60 and GRP75 under basal as well as paraquat-induced stress conditions. Even though human SLP-2 has been shown to interact with CL-containing liposomes, the affinity towards PA has not been evaluated so far. Therefore, we investigated whether purified human SLP-2 can directly bind to PA, like it’s *C. elegans* counterpart STL-1. We performed the lipid overlay assay for human SLP-2, which was purified as a recombinant His-tagged SLP-2, expressed in insect cells (fig. S6, A to C). Using the lipid membrane strip, we determined that human His-SLP-2 also demonstrates the highest binding affinity to PA, and to a lesser extent to PS, CL, and sulfatide (Fig. 6, C and D). These data thus reveal an evolutionarily conserved role of SLP-2 in PA binding and the mtUPR.

**Fig. 6.**
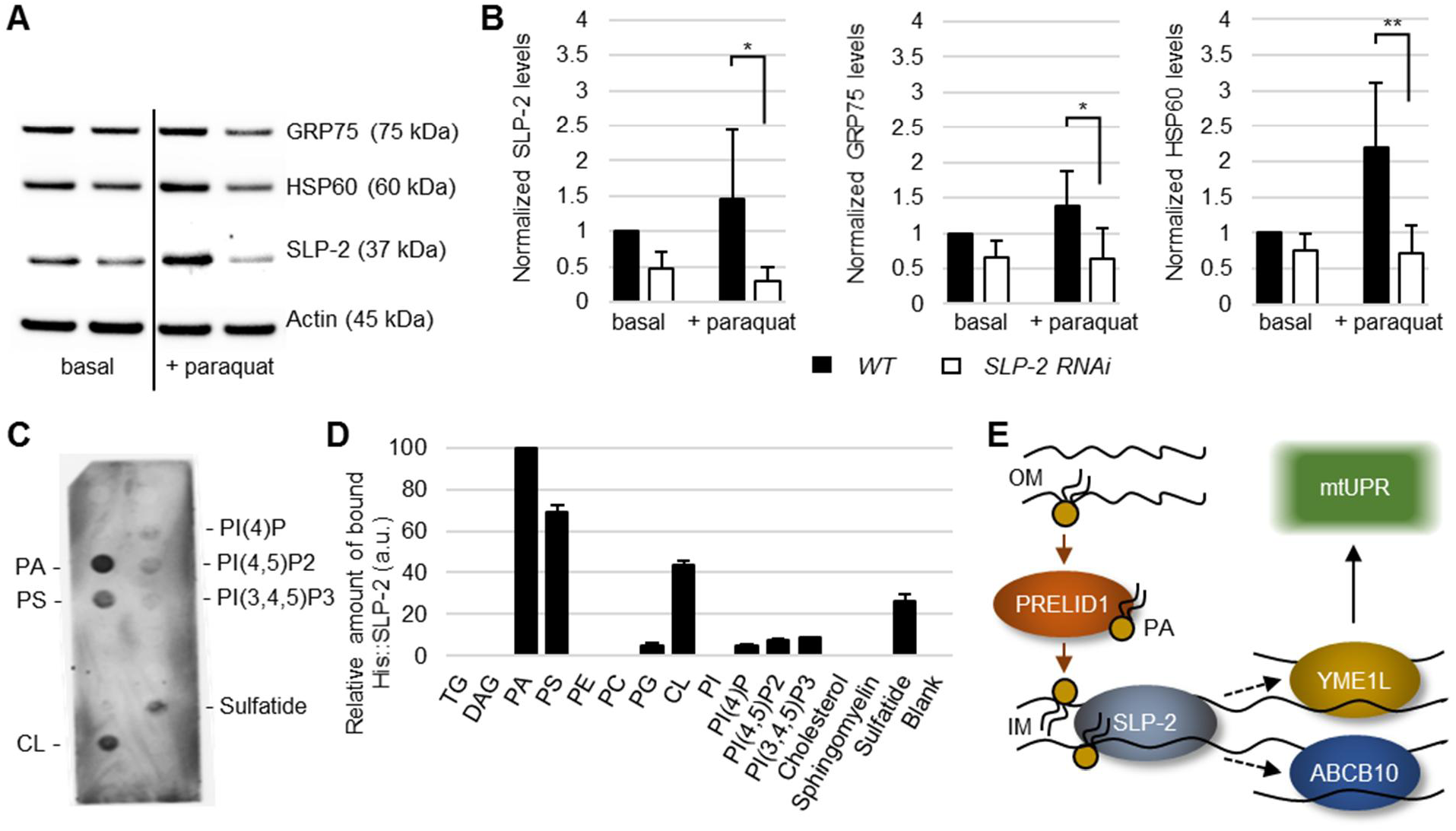
Conserved role of SLP-2 in the mtUPR and PA binding. **A.** Western Blot analysis of human mtUPR targets using whole cell extracts of mammalian SH-SY5Y neuroblastoma cells. **B.** Semi-quantification of the respective protein signal from the Western Blot analysis on the left. Three independent biological replicates, **P< 0.01, *P< 0.05. **C.** Lipid overlay assay using purified SLP-2::His followed by anti-SLP-2 serum. PA; Phosphatidic acid, PS; Phosphatidylserine, CL; Cardiolipin **D.** Semi-quantification of bound His::SLP-2 on indicated lipid spots with three technical replicates. **E.** Proposed model of the SLP-2-mediated mtUPR upon PA binding.

## DISCUSSION

The protein SLP-2 plays an important role in mitochondrial biology during oxidative stress, such as stabilization of several proteins located within mitochondrial membranes, or stimulation of CL biosynthesis or its stability (32). However, its exact mode of action is unclear. In this study, we propose a model in which SLP-2 senses mitochondrial fitness through physical interaction with PA at the inner mitochondrial membrane to promote the mtUPR, and thereby confer mitochondrial protection during oxidative stress (Fig. 6E).

Our data indicate that *stl-1* is not required for the mtUPR when mitochondria are damaged upon severe stress, but rather plays a regulatory role in stress detection via sensing of lipid remodelling in the mitochondrial membrane, enough to mediate sufficient mtUPR activation when needed. A previously reported functional, as well as physical connection between SLP-2 and CL may suggest their mutual importance in the mtUPR (33). In fact, the role of CL in regulation of the mtUPR is mediated through the stabilization of several mitochondrial protein complexes in the inner mitochondrial membrane, including the protein import machinery (34). However, our data support a model in which PA, but not CL, is the key element for STL-1/SLP-2-mediated mtUPR. First, we found a discrepancy between initial (*prel-1*) and later (*pgs-1, crls-1, acl-3*) steps of CL biosynthesis in the activation of the mtUPR. Specifically, nematodes with reduced CL levels induce the mtUPR in an *stl-1*-dependent manner, but not those worms with disrupted PA homeostasis at the inner mitochondrial membrane. In line with this finding, inactivation of both *Ups1* (ortholog of human PRELID1) and *Crd1* (ortholog of human *CRLS1*) genes results in CL depletion in a yeast model. However, only loss of yeast *Ups1* impairs the basal mtUPR indicating that a PA-mediated mtUPR is an evolutionary conserved mechanism conferring mitochondrial protection (26,35,36). Second, we show that both purified STL-1 and SLP-2 physically interact with PA in a pH-dependent manner. The pKa (acid dissociation constant) of PA and CL has been determined around pH 6.9-7.9 and pH 7.5-9.5, respectively, depending on the environment, making PA a more suitable lipid species for sensing oxidative stress (37–39). Intriguingly, elevation of the paraquat concentration makes the thylakoid membranes more acidic (40), supporting a notion that the STL-1 interaction with PA upon oxidative stress can be driven by a change in pH due to disrupted proton transport in mitochondria. Another possibility of PA sensing by STL-1 can be mediated by the local lipid concentration changes, as supported by the fact that inhibited CL biosynthesis, which results in an increased ratio of PA/CL, activates the mtUPR in an *stl-1*-dependent manner. Third, using NAO staining we show that oxidative stress induced by paraquat changes lipid composition of the mitochondrial membrane, in a way that resembles the phenotype observed upon disruption of CL biosynthesis. While abundant mitochondrial CL species are prone to oxidation/peroxidation (41), PA has a minor representation in the mitochondrial membrane, yet its level in mitochondria increases during oxidative stress and plays a role in mitochondrial fusion (42,43). Reduced CL levels, either through peroxidation or inhibited biosynthesis, thus dramatically alter the PA/CL ratio, which very likely alters the STL-1 structural organisation and/or localization as demonstrated by an increased formation of bright fluorescent foci of STL-1::GFP in CL-depleted mitochondria upon stress. Based on these data, we conclude that the physical interaction between PA and STL-1 regulates the ability of STL-1 to sustain mtUPR stimulation under basal conditions and mediate/trigger a stronger mtUPR when oxidative stress increases. Nonetheless, further studies are needed to delineate the exact mechanism of PA-mediated regulation of STL-1/SLP-2. In particular, the role of other phospholipids in SLP-2 regulation and the downstream effect of SLP-2-lipid interactions, such as plausible structural rearrangement of SLP-2 and regulation of the SLP-2-protein heterocomplexes, merits further investigation.

Based on our data, we can speculate that SLP-2 regulates the mtUPR through a functional interaction with YME1L; both human SLP-2 and YME1L are subunits of the SPY complex that regulates the processing and degradation of various proteins in the mitochondrial intermembrane space (13), both proteins are activated in stress conditions (44,45), and inactivation of both *C. elegans* orthologs, *stl-1* and *ymel-1*, leads to compromised mtUPR. Moreover, it has been shown that the alteration of the mitochondrial lipid composition upon stress signalling leads to proteolytic rewiring of mitochondria by YME1L (45). Therefore, we speculate that *stl-1* together with *ymel-1* may constitute the mtUPR activation pathway through intermembrane peptide-mediated signalling. In fact, specific proteolytic processing of mitochondrial proteins by the YME1L machinery initiates various downstream signalling pathways suggesting that protein processing might generate signals for the mtUPR activation (46). As a target of YME1L activity, PREL-1/PRELID1 might play a dual role in mtUPR activation, either by transporting PA into inner mitochondrial membranes and/or by providing signalling peptides through its processing. Interestingly, suppression of the mtUPR in *stl-1* and *ymel-1* mutants resembles matrix peptide-mediated suppression of the mtUPR in mutants with inactivated *haf-1* that encodes the mitochondrial inner-membrane transporter ABCB10 (30). Human ABCB10 is involved in the mammalian mtUPR similarly to SLP-2 as demonstrated in this study. Thus, STL-1/SLP-2 regulating HAF-1/ABCB10 gives rise to the possibility of a downstream *stl-1* signalling pathway for mtUPR activation. However, the possible association between *prel-1*, *stl-1*, *ymel-1,* and *haf-1* to mediate the mtUPR also needs to be addressed in future studies.

The PA binding activity of SLP-2 might also explain the beneficial role of both PA as well as SLP-2 in stress through CL biosynthesis as well as regulation of mitochondrial dynamics. It is plausible that SLP-2 organizes PA microdomains in the inner membrane to spatially regulate CL biosynthesis or stability altering inner membrane lipid composition for fusion/fission (11,15,16,42). In addition, SLP-2 also interacts with, and stabilizes, Mitofusin (MFN) (14), an outer mitochondrial membrane factor mediating fusion of the mitochondrial outer membrane, an event regulated by PA (42), and which, through YME1L activity, stabilizes OPA1 (13,16,45), a dynamin-like GTPase responsible for fusion of the inner mitochondrial membrane.

Together, our results implicate two possible outcomes of PA-bound SLP-2: (1) PA regulates SLP-2 specificity for its protein interacting partners, such as YME1L and/or ABCB10, and (2) SLP-2 forms PA microdomains for targeted CL biosynthesis. A recent study reported that the closest ortholog of SLP-2 in *Trypanosoma brucei* TbSlp2 can also bind to PA *in vitro* (47) suggesting that SLP-2/PA-mediated mtUPR is a fundamental mechanism of mitochondrial protection, which is evolutionary conserved from unicellular organisms to humans. A direct involvement of SLP-2 in the processing of the Parkinson’s disease related protein PINK1 by YME1L, together with the fact that mild overexpression of SLP-2 restores mitochondrial impairment in Parkinson’s disease cellular and *Drosophila* models, reveals a therapeutic potential of SLP-2 (12,13,48,49). A regulatory role for mitochondrial PA through SLP-2 thus provides new possibilities of therapeutic targeting for diseases with mitochondrial involvement.

## MATERIALS AND METHODS

### Caenorhabditis elegans Strains

Animals were maintained by standard procedures on nematode growth media (NGM) plates at 20°C. The strains used were as follows: N2, *zcIs9(hsp-60p::GFP), zcIs13(hsp-6p::GFP), eraIs4(stl-1p::stl-1::GFP), stl-1(tm1544), stl-1(gk118866), stl-1(gk118865), phb-2(ad2154), eat-3(ad426), fzo-1(tm1133), drp-1(tm1108), ymel-1(tm1920), haf-1(ok705), aco-2(rf40[aco-2::GFP]), foxSi75([eft-3p::tomm-20::mKate2::HA::tbb-2 3’ UTR)*.

### Transgenesis

*C. elegans* constructs for the *eraIs4* transgene were generated by direct PCR of the entire *stl-1* gene using primer stl-1_A, stl-1_B (Table S1), and subsequently fused in-frame with the *GFP* reporter gene with *unc-54 3′*UTR which was amplified from pPD95.75 plasmid (Addgene) using GFP_C and GFP_D primers (Table S1). The PCR product was injected into Bristol strain N2, and an extrachromosomal array was established. The extrachromosomal array was subsequently integrated by ultraviolet (UV) irradiation, and the animals carrying *eraIs4* were subjected to 5× outcrossing.

### Paraquat treatment

100 µl of OP50 bacterial culture containing 1mM paraquat was spread onto plates containing 5 ml of solid NGM. Plates were air-dried overnight before usage in the assay. Worms were placed onto the plates containing paraquat for the indicated time before being evaluated. SH-SY5Y cells were incubated in medium containing 1mM paraquat for 24 hours before being evaluated.

### 10-N-nonyl acridine orange (NAO) staining

100 µl containing 100µM NAO in M9 buffer (22mM KH_2_PO_4_, 42mM Na_2_HPO_4_, 85.5mM NaCl, 1mM MgSO_4_) was spread onto NGM plates with nematodes crawling on the bacterial lawn. Nematodes were incubated on the plates containing NAO for 2 hours before being analysed by confocal microscopy with excitation and emission wavelengths set at 490 and 510-525 nm, respectively.

### RNA interference

Feeding RNAi was performed as previously described. Five gravid animals were placed on NGM media containing ampicillin 25 μg/ml and 1mM Isopropyl β-d-1-thiogalactopyranoside (IPTG) and seeded with bacteria producing the desired double stranded RNA (dsRNA). Progeny were subsequently grown at 20°C and screened for the phenotype in young adult stage. Visual examination of the animals was done under the fluorescent stereoscope (Nikon SMZ800N). The bacterial clones were obtained from *C. elegans* RNAi collection - Ahringer (Source Bioscience, 3318).

### Determination of GFP levels

One-day-old adult hermaphrodites were lysed in the Laemmli sample buffer (Bio-Rad) supplemented with the reducing agent β-mercaptoethanol, followed by boiling for 10 min. The worm lysates were separated on 4–15% sodium dodecyl sulphate-polyacrylamide gel electrophoresis (SDS PAGE) (Bio-Rad). The proteins were transferred to a nitrocellulose membrane (Bio-Rad) and subsequently detected by anti-GFP rabbit polyclonal antibody (Novus Biologicals #NB600-308; 1:1000) and re-incubated with anti-Actin rabbit polyclonal antibody (Cell Signaling #4967; 1:2000) as a loading control. Western blot signals were semi-quantified in ImageJ software (Fiji).

### Quantitative real-time polymerase chain reaction analysis

Grown plate of respective *C. elegans* strain of mixed developmental stages was collected and washed in M9 buffer. The nematode pellet went 2x through the freeze/thaw cycle before sonication. Total RNA was extracted using the Direct-zol RNA Miniprep kit (Zymo research) using TRIzol® Reagent (Thermo Fisher Scientific). 1 µg of RNA was retrotranscribed using the SuperScript VILO cDNA Synthesis Kit (Thermo Fisher Scientific). The qRT-PCR was performed on the CFX96 Real-Time PCR Detection System (Bio-Rad) using All-in-One SYBR® Green qPCR Mix (GeneCopoeia) with specific pairs of primers for *hsp-60*, *hsp-6*, and *act-2* as a reference, using the following conditions: 95 °C for 10 mins; 40 cycles of 95 °C for 10 sec; 58 °C for 20 sec; 72 °C 15 sec. Melting curve analysis: 72° to 95 °C, increment 0.5 °C, for 6 sec. Primer sequences are listed in Table S1.

### Purification of STL-1::GFP

Animals *eraIs4* expressing STL-1::GFP were collected and washed in M9 buffer. The nematode pellet went 2x through the freeze/thaw cycle before sonication in M9 buffer containing 0.1% Triton X-100 and protease/phosphatase inhibitor mixture (Sigma-Aldrich). Whole-animal extract was centrifugation-cleared for 30 mins at 4°C, and STL-1::GFP was captured by immunoprecipitation with GFP-trap magnetic beads (ChromoTek) at room temperature for one hour. The GFP-trap beads were subsequently washed five times with M9 buffer containing 0.1% Triton X-100. Co-immunoprecipitated proteins were eluted by glycine elution buffer pH 2.5 and immediately neutralized by the neutralization buffer. Purification fractions and eluates were then subjected to SDS-PAGE or immediately used in lipid overlay assays, respectively.

### Expression and purification of recombinant His::STL-1

25 ml of LB media containing Kanamycin (50ug/ml) in the 50 ml Falcon tube was inoculated with BL21(DE3) expressing cells carrying pET-28a with cloned *C. elegans stl-1*. The *stl-1* was amplified without the part encoding the mitochondrial signal sequence using primers STL-1_His5 and STL-1_His3, and subsequently cloned between *NdeI* and *XhoI* restriction sites ensuring production of His tagged STL-1 at the N-terminus. Bacteria carrying *pET28a-stl-1* were shaken for 16 hours at 25°C to high optical density, and subsequently protein expression was induced by adding IPTG into 0.1mM final concentration. Bacterial cultures were shaken at 25°C for another 3 hours before being collected by centrifugation at 4°C for 10 minutes. Bacterial pellet was washed in 1xPBS and centrifuged at 4°C for 10 minutes. Lysis buffer (20 mM HEPES, 500 mM NaCl, 20 mM imidazole, 10 mM BME, 0,1 % Triton X-100 and protease inhibitor cocktail) was used to solubilize the bacterial pellet followed by sonication until the lysate was clear. Bacterial lysate was centrifuged at 20,000g for 30 minutes and supernatant was collected. The His::STL-1 was purified using Nuvia IMAC column (Bio-Rad) in the presence of washing buffer (20 mM HEPES, 500 mM NaCl, 20 mM imidazole, 10 mM BME, 0,1 % Triton X-100). Soluble fraction of the lysate (supernatant) was loaded onto the Nickel column using peristaltic pump. The column with bound His-tagged proteins was washed in washing buffer (20mM HEPES, 500mM NaCl, 20mM imidazole, 10mM BME, 0,1 % Triton X-100) and then eluted by an imidazole gradient from 20 to 500mM over 20 column volumes. During the purification in the presence of 1% CHAPS, 0.1% Triton X-100 was omitted in the buffers.

### Size exclusion chromatography

Size exclusion chromatography (SEC) was used to determine particle size of the purified STL-1 incorporated into the detergent micelles. The SEC was performed in 20 mM HEPES, 500 mM NaCl, 10 mM BME and either 0.1% Triton X-100, 1% CHAPS or without any detergent using Enrich 650 column (BioRad). The column was calibrated with APOA-4 composed of monomeric (40 kDa) and dimeric (80 kDa) species.

### Expression and purification of SLP-2::His

The protein was expressed in Sf9 insect cells from infection with P2 virus. Cells were harvested by centrifugation at 4°C, and the cell pellet was collected and washed with 1x PBS and snap freezing in liquid nitrogen before being sonicated at amplitude. The extract was clarified by centrifugation at 18,000 x g for 1 hour. Nickel beads were washed and equilibrated in base buffer (50mM HEPES pH 7.5, 500mM NaCl, 10% glycerol, 10mM imidazole, 1mM TCEP), subsequently transferred to the supernatant, and incubated for one hour in the cold-room with gentle end-over-end rotation. The supernatant was transferred to the clean Econo column. Collected beads were washed with washing buffer (50mM HEPES pH 7.5, 500mM NaCl, 10% glycerol, 45mM imidazole, 1mM TCEP). Elution was performed by applying Elution buffer (50mM HEPES pH 7.5, 500mM NaCl, 10% glycerol, 350mM imidazole, 1mM TCEP) to the beads and incubating for 5 min each time for each elution fraction. Samples were additionally injected into the Ӓkta system, and gel filtration was run in Buffer-GF:50mM HEPES pH 7.5, 0.2M NaCl, 5% glycerol and 1mM TCEP. The obtained fractions were analysed by SDS-PAGE and collected for further analysis.

### Lipid overlay assay

The membrane lipid strip was air-dried, blocked with 3% protease-free BSA in Tris-buffered saline with 0.1% Tween (TBST) for 1 h, overlaid with either STL-1::GFP, His::STL-1 or His::SLP-2 protein (0.5 μg/ml in blocking buffer) for 1 h. After three washes with 1x TBST, the membrane was incubated with primary rabbit anti-GFP, rabbit anti-His Tag or mouse anti-SLP-2 antibody, respectively, for 1 h or overnight at 4°C in blocking buffer, followed by three washes with TBST and incubating the membrane with secondary antibody for chemiluminescence imaging.

### Imaging

Animals were mounted on a 2% agarose pad with 10 mM sodium azide and imaged on a Leica SP8-X confocal laser scanning microscope within 5–10 min. To study the expression pattern of STL-1::GFP and NAO staining, we used polystyrene beads to avoid anaesthetics. At least three images representing each *C. elegans* strain from three independent biological replicates were analysed. Colocalization analysis was done using the colocalization threshold plugin interface in the ImageJ application followed by measurement of colocalized vs non-colocalized area within dual colour image. The GFP (Venus) fluorescence was imaged with 489 nm excitation and 500-550 nm emission range, mKate2 with 589 nm excitation and 600-650 nm emission, and NAO staining with excitation at 490 nm and 510-525 nm emission. At least three images representing each *C. elegans* strain from three independent biological replicates were analysed.

### Cell models

Human neuroblastoma SH-SY5Y cells (ATCC CRL-2266) were cultured in Dulbecco’s modified Eagle’s Medium (DMEM, Sigma) supplemented with 10% fetal bovine serum and 1% penicillin-streptomycin (Thermo Fisher Scientific). SH-SY5Y cells with a stable knockdown of SLP-2 were produced as described previously (12). Cells were maintained at 37 °C in a saturated humidity atmosphere containing 5% CO2. Whole cell lysates of SH-SY5Y cells (10 µg) were subjected to Western blotting using the following antibodies: mouse anti-SLP-2 antibody (Abcam ab89025, 1:1000), rabbit anti-HSP60 (Cell Signaling #4870; 1:1000), rabbit anti-GRP75 (Abcam, ab53098; 1:5000), and rabbit anti-β actin (Cell Signaling. #4967; 1:1000).

### Statistical Analyses

Data are presented as mean ± standard deviation (SD). Statistical significance was determined using one-way analysis of variance (ANOVA) with Holm correction for multiple comparisons and the unpaired two-tailed *t* test for single comparisons. The threshold for significance was set at p < 0.05.

## ACKNOWLEDGEMENTS

We thank the Caenorhabditis Genetics Center, which is funded by NIH Office of Research Infrastructure Programs (P40 OD010440) and National BioResource Project (NBRP)::C. elegans (Mitani lab) for some *C. elegans* strains, and Dengke Ma lab for some plasmids. Furthermore, we thank Dr. Nicola Burgess-Brown, Dr. Alejandra Fernandez, and Dr. Leslye Roca Burgos from the CMD, University of Oxford (UK) for the production of recombinant human SLP-2.

## Author contributions

M.P.C.R., K.M., A.H. and R.V. performed the experiments, M.P.C.R., I.P., A.A.H., A.H. and R.V. designed the experiments, P.C., K.M., I.P., A.A.H., A.H., A.L. and R.V. analysed the data, I.P, S.K., P.P.P., A.A.H., A.H. and R.V. acquired funding, I.P., S.K., A.A.H. and A.H. helped to revise and edit the manuscript, R.V. wrote the manuscript. All authors approved the final version of the manuscript.

## Data and materials availability

All data needed to evaluate the conclusions in the paper are included in the paper and Supplementary material. Further inquiries can be directed to the corresponding author.

## SUPPLEMENTARY MATERIAL

Figs S1 to S6

Table S1

